# Serial dependence in the perception of proportion

**DOI:** 10.1101/2025.10.29.685242

**Authors:** Kaede Hashiguchi, Yuichi Tanaka, Hiroshi Higashi

## Abstract

The influence of previously encountered stimuli on subsequent perceptual judgments is termed serial dependence. Serial dependence has been robustly demonstrated across a broad spectrum of visual features, ranging from low-level attributes, such as orientation and color, to high-level representations, including facial identity. To further understand this dependency, the present study investigated the presence of serial dependence in judgments of relative proportions within visual stimuli. Participants completed a two-alternative forced-choice (2AFC) task involving stimuli composed of two symbols. These symbols differed only in brightness in Experiment 1 and only in shape in Experiment 2. Participants were required to judge which symbol occupied the larger proportion. The results revealed a systematic bias in the participants’ responses. Specifically, Experiment 1 yielded a repulsive bias (negative serial dependence), where current judgments shifted away from the proportion observed in the preceding trial. Conversely, Experiment 2 exhibited an attractive bias (positive serial dependence), pulling current judgments toward the preceding proportion. Furthermore, the magnitude and direction of this serial dependence appeared to be modulated by both the participants’ confidence in their judgments and the feature similarity between successive stimuli. These findings suggest that the perception of proportion has serial dependence and extend the generality of this phenomenon to higher-level visual summary statistics.

## 1 Introduction

Retinal images, formed by focusing incoming light onto the retina, are inherently unstable, constantly fluctuating due to both external and internal noise (e.g., changes in ambient lighting, eye blinks). Despite this sensory variability, our perception of the world remains remarkably stable; objects do not appear to jitter or shift unpredictably from moment to moment. This striking perceptual constancy is hypothesized to arise because the visual system leverages the continuity of the visual world, utilizing recently encountered information to mitigate sensory noise [1–4]. This compensatory mechanism is behaviorally evidenced as serial dependence in visual judgments [1, 5]. Serial dependence is defined as a behavioral bias where current visual judgments are systematically influenced by previously seen stimuli [5]. For instance, Fischer and Whitney [5] demonstrated this effect by presenting a tilting Gabor stimulus and asking participants to report its orientation. Their results showed that adjustment errors positively correlated with the orientation difference between the current and preceding stimuli. This finding indicates an attractive bias―responses are pulled toward the orientation of the preceding stimulus―a phenomenon termed positive serial dependence. Crucially, the magnitude of this bias became greater when the current and previous orientations were similar, diminishing or disappearing as the difference increased. While the effect decayed over time, serial dependence persisted for stimuli presented up to three trials earlier (approximately 10–15 seconds).

Further studies have established that serial dependence is a generalized phenomenon, affecting a vast array of visual features beyond simple orientation [6–10]. These features span from low-level properties―including spatial position [11–13], color [14–16], numerosity [17–20], and shape [21,22]― to high-level, complex features such as face identity [23–26], facial expression [27,28], and perceived age [29, 30].

While these studies have fundamentally advanced our knowledge of the mechanisms and ubiquity of serial dependence, our understanding remains incomplete. To fully delineate the conditions and underlying mechanisms of this bias, it is essential to extend investigations to perceptual visual features that have not yet been examined [31].

The current investigation seeks to determine whether serial dependence occurs in the perception of proportion within visual stimuli [32]. The perception of proportion is a vital aspect of ensemble perception―the ability to rapidly estimate the relative quantity of components in a visual array. Examples include judging whether “about half” of a group of items share a feature (e.g., red apples in a basket) or quickly noting that “most” seats in a room are occupied. Such perceptual estimates allow for the rapid extraction of overall summary statistics without the need for painstaking individual element enumeration.

We conducted two behavioral experiments using arrays of symbols as visual stimuli. Experiment 1 focused on brightness-based perception of proportion. A stimulus was composed of 32 symbols, identical in size and shape but varying in brightness across two predefined levels. Experiment 2 then sought to determine whether any observed serial dependence was specific to brightness or a more general property by varying the proportion for symbols of two shapes.

In both experiments, participants performed a two-alternative forced-choice (2AFC) task, judging which of the two symbol types (brightness in Experiment 1; shape in Experiment 2) occupied a greater proportion in the display. To quantify the serial dependence, we examined the Point of Subjective Equality (PSE), defined as the proportion at which the participant perceived the two symbols as equally represented. The presence of serial dependence was evaluated by examining whether the proportion of the two symbols in the preceding trial contributed to variations in the PSE.

The findings of this study support the existence of serial dependence in the perception of proportion within visual stimuli. Furthermore, our results showed that serial dependence increased when the proportion of the two symbols in the preceding trial was easier to discriminate. In addition, changes in the direction of serial dependence appeared to be influenced by the degree of similarity between the stimuli of the preceding and current trials. These findings extend previous research by demonstrating that both the stimulus discriminability (a proxy for participant confidence) and the inter-trial feature similarity are key modulators of serial dependence in the perception of proportion.

## 2 Materials and methods

### 2.1 Participants

A total of 200 participants were recruited for this study via the online platform Prolific [33], with data collection conducted using Pavlovia.org [34]. One hundred participants were enrolled in each experiment. In Experiment 1, the sample (*N* = 100) included 50 female participants, with a mean age of 34 years (range: 18–67 years). Similarly, Experiment 2 enrolled 100 participants (50 female), with a mean age of 34 years (range: 20–67 years). This study was approved by the Committee for Human Research at the Graduate School of Engineering, The University of Osaka (Approval number: 5-4-1), and complied with the Declaration of Helsinki. Prior to participation, all individuals were provided with a detailed explanation of the experimental procedures and the associated monetary compensation, and informed consent was obtained before they began the study. Each participant received 3 GBP as compensation. For all participants, the following information were recorded: sex, age, simplified ethnicity, country of birth, country of residence, nationality, language, student status, employment status, and screen characteristics.

### 2.2 Stimuli

Stimuli were generated using PsychoPy software (v.2024.1.5). The experiment was performed on an online platform and conducted through a web browser on each participant’s personal computer. Consequently, it was not possible to exert precise control over the position and size of the stimuli. Therefore, the position and size of the stimuli were specified relative to the height of the participant’s browser window. To prevent the stimuli from appearing too small, participants were required to maintain full-screen mode throughout the experiment.

The stimulus display is illustrated in Figure 1. All visual stimuli were presented against a uniform gray background (RGB: 128, 128, 128) in the standard sRGB color space. Let *h* denote the height of the participant’s browser window. A fixation cross (white; RGB: 255, 255, 255), consisting of vertical and horizontal lines, was presented at the center of the screen with a height of 0.02*h*. This cross served to maintain participants’ central gaze during inter-stimulus intervals. The test stimulus consisted of 32 individual symbols, each having a height of 0.02*h*. These symbols were pseudo-randomly positioned within a circular display area centered on the screen, limited to a radius of 0.2*h*. Crucially, the minimum distance between the centers of any two symbols was constrained to be greater than or equal to 0.02*h*. The exact spatial positions were randomized on every trial.

**Figure 1:**
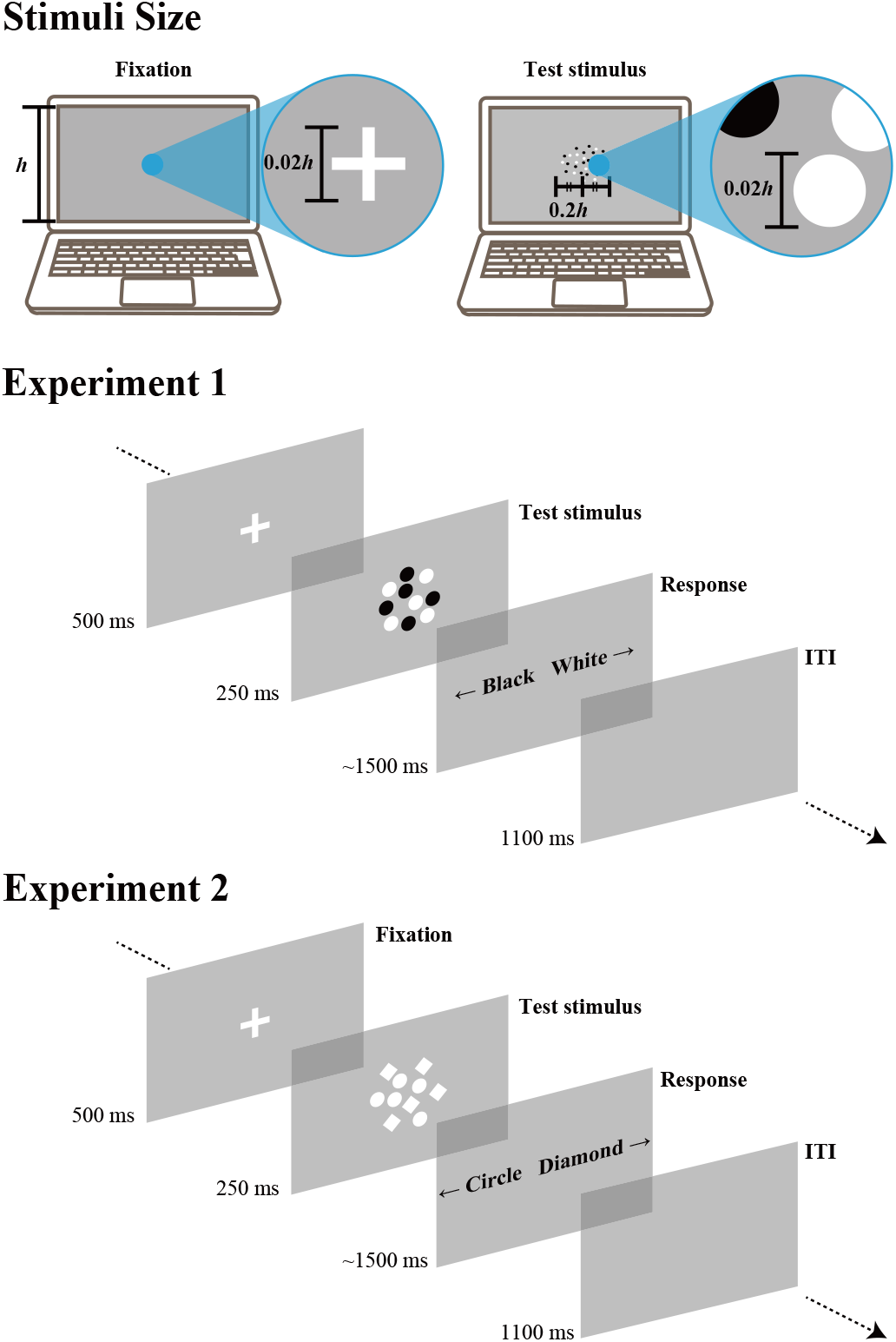
Stimulus sizes and general experimental procedure. In both experiments, the height of the white cross was set to 0.02 times the height of the browser window. The height of each symbol in the test stimulus was also 0.02 times the browser window height. All symbols were presented within a range of 0.2 times the browser window height from the screen center. In Experiment 1, a white fixation cross was first presented at the center of the screen for 500 ms. This was followed by a test stimulus composed of white and black dots, presented for 250 ms. The proportion of white and black dots varied across trials. Participants were asked to indicate, in a 2AFC task, which brightness―white or black―occupied a larger proportion of the stimulus. After a response was made or 1500 ms had passed, a blank screen was shown for 1100 ms before the next trial began. In Experiment 2, the test stimulus was composed of dots and diamonds, and participants indicated which shape was more prevalent in the stimulus.

The two experiments employed distinct features to manipulate proportion. Experiment 1 (Brightness Proportion) used a stimulus array composed of circular dots that differed only in brightness: white (RGB: 255, 255, 255) and black (RGB: 0, 0, 0). On each trial, the target proportion of white dots was randomly selected from the set {10%, 40%, 45%, 50%, 55%, 60%, 90%}, with the remainder being black. The corresponding number of white dots for each proportion was rounded to the nearest integer. Experiment 2 (Shape Proportion) employed an array comprising two different shapes that were equated for brightness (both white): circular dots and diamond shapes. The diamond was defined as a square rotated by 45°, with a diagonal length of 0.02*h* (thus matching the dot’s height). The target proportion of dots versus diamonds was selected using the identical set of proportions used in Experiment 1.

### 2.3 Procedure

The entire experimental session consisted of two sequential blocks: a practice block followed by the main test block.

The procedure for a single trial in the test block of Experiment 1 is depicted in Figure 1. Each trial commenced with the presentation of a white fixation cross for 500 ms, and participants were instructed to maintain fixation throughout its display. This was immediately followed by the 250 ms presentation of the test stimulus, which was composed of white and black dots. After the stimulus disappeared, participants were prompted to report which brightness of dot―white or black―constituted the greater overall proportion in the set. Responses were registered by pressing the left or right arrow key. The key-to-brightness mapping was randomized and counterbalanced across participants, but remained fixed for each individual throughout the experiment. Participants given a 1500 ms limit from the disappearance of the test stimulus to respond, and were instructed to respond as quickly as possible. Failure to respond within this time limit was recorded as a no-response. Following a valid response or the expiration of the 1500 ms limit, a 1100 ms blank screen preceded the next trial. Participants received a one-minute break upon completion of half the trials. The test block consisted of 420 trials and the entire experiment took approximately twenty minutes to complete.

The procedure for Experiment 2 was identical to that of Experiment 1, with the critical exception being the nature of the test stimulus. In Experiment 2, the stimulus was composed of white dots and white diamonds, and participants were required to judge the greater shape proportion (dots vs. diamonds). The key-to-shape mappings were similarly randomized and counterbalanced across participants.

Prior to the main test block in both experiments, participants completed a practice block consisting of two trials before beginning the main test block of 420 trials. The stimulus presentation and response procedures in the practice block mirrored those of the test block. However, in the practice trials, the proportions of the target stimulus were fixed at 20% and 80% (order randomized), and participants received feedback. If a response was made within the 1500 ms limit, they were informed whether their answer was correct or incorrect; if they failed to respond in time, they were notified that their response was too slow. This practice block was designed to familiarize participants with the experimental flow, train them on the key-response correspondence, and establish the time constraints, thereby promoting accurate and speedy responses during the subsequent test block.

### 2.4 Behavioral data analysis

Behavioral data underwent an initial two-stage screening process to ensure participant focus and reliability. First, to exclude participants who failed to remain engaged with the task and had an excessive number of no-responses within the time limit, the no-response rate (*r*_nr_) was calculated for each participant. Outliers were identified using the interquartile range (IQR) method: the first quartile (*Q*_1,nr_) and the third quartile (*Q*_3,nr_) of the no-response rates were computed across participants, defining the interquartile range as IQR_nr_ = *Q*_3,nr_ − *Q*_1,nr_. Participants whose noresponse rate exceeded the upper boundary defined by the following criterion were excluded.

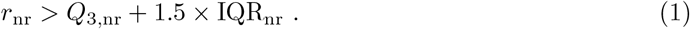

Next, an accuracy-based screening was applied to the remaining dataset to exclude participants who responded but appeared to be inattentive―potentially responding randomly. Accuracy was calculated for each participant, focusing exclusively on trials with highly skewed proportions (10% or 90% white dots). The reason for limiting the accuracy calculation to these two stimulus conditions was to minimize variability in accuracy caused by the influence of serial dependence. When serial dependence is present, participants’ responses can be biased by the stimulus shown in the preceding trial. This influence may lead to an increased rate of incorrect responses. Consequently, if accuracy were calculated across all trials, it would become difficult to determine whether variability in participants’ accuracy was due to random responses or the influence of serial dependence. Therefore, we focused on trials with highly skewed proportions, in which the correct answer was obvious. By focusing on trials where the correct answer was highly obvious, we could more accurately identify participants who were inattentive. As in the previous step, the first quartile (*Q*_1,acc_) and third quartile (*Q*_3,acc_) of the accuracy rates (*r*_acc_) for these targeted trials were computed. Participants whose accuracy rate fell below the lower boundary were excluded from subsequent analysis:

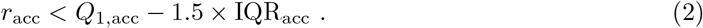

As a result, we excluded 17 participants from data analysis in Experiment 1 and 22 participants in Experiment 2.

For the remaining valid data, trials were subsequently organized into four subsets for each participant based on the proportion of the white dots presented in the immediately preceding trial: 10%, 40%, 60%, or 90%. Preceding trials with proportions of 45%, 50%, or 55% were excluded from this subset analysis. This exclusion was made because the near-parity proportions in these stimuli made it difficult to determine the dominant symbol, thus reducing the likelihood that they would exert a robust biasing effect on the perception of the subsequent stimulus. A psychometric function was then fitted to the response data within each subset using the cumulative distribution function:

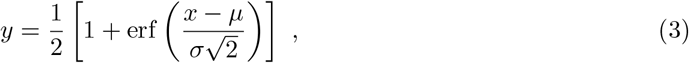

where *y* is the response rate for each proportion of white dots, *x* is the proportion of white dots, and *µ* and *σ* are the mean and scale of the cumulative distribution function, respectively. The function erf(·) denotes the error function, which is defined by

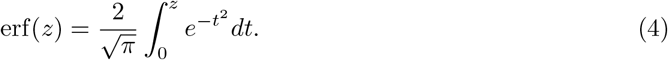

The PSE, reflecting the accuracy of each participant’s proportion discrimination performance, was defined as the value of *x* at *y* = 50. The PSE value varies across subsets depending on the presence and magnitude of serial dependence.

To quantitatively evaluate the temporal evolution of PSE differences across subsets caused by serial dependence, we calculated the following the PSE differences for each participant:

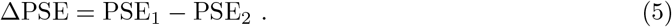

Here, ΔPSE represents the difference in the PSE between subsets, PSE_1_ and PSE_2_ refer to the PSE for any two subsets. After calculating ΔPSE for each participant, we then computed the average across all participants. To examine whether there was a significant differences in the PSE across subsets, a one-sample t-test was conducted to determine whether the null hypothesis―that the mean difference is zero―could be rejected.

## 3 Results

### 3.1 Experiment 1 (White vs Black)

Because the present study employed a 2AFC task, it was necessary to rule out the possibility that the observed PSE shifts were driven by non-perceptual response strategies, such as an unconscious tendency to repeat or alternate responses when uncertain. Such behavior could lead to shifts in the PSE and potentially influence the evaluation of serial dependence. To address this, we calculated the proportion of trials in which each participant’s current response matched their previous response. A proportion exceeding 50% would indicate a repetition tendency, while a proportion below 50% would suggest an alternation tendency. The analysis showed no systematic tendency toward either repetition or alternation for either the right key (*t*(82) = −1.21, *p* = 0.23) or the left key (*t*(82) = −0.94, *p* = 0.35), as illustrated in Figure 2A. This finding suggests that the observed PSE differences reflect perceptual effects related to stimulus processing rather than non-specific response patterns.

**Figure 2:**
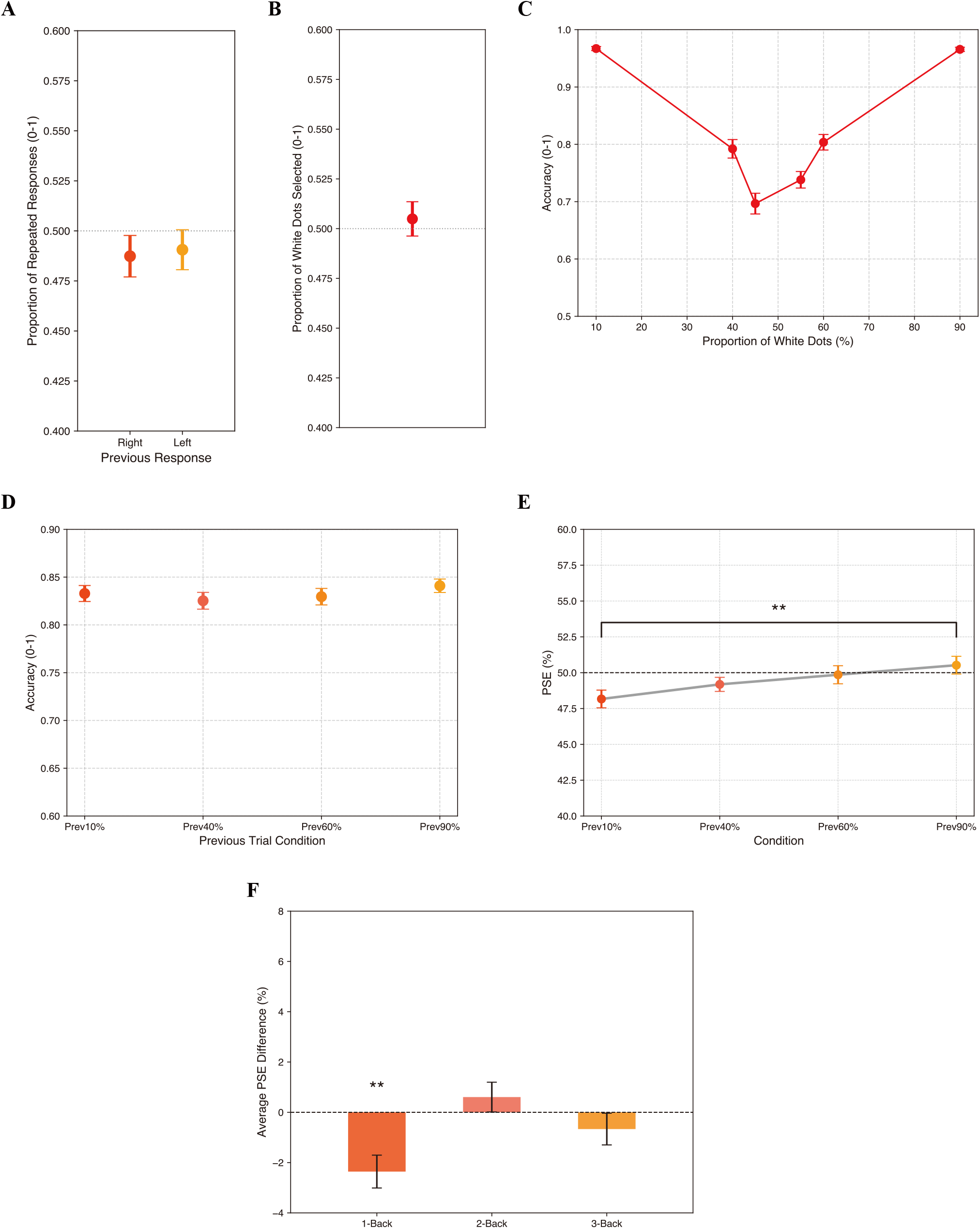
Results from Experiment 1. Error bars in (A)–(F) represent the standard error of the mean across participants. (A) Probability of repeating the same response as the previous trial. (B) The mean proportion of “more white dots” responses across all participants. (C) Overall accuracy across participants for each proportion of white dots. (D) Overall accuracy across participants for each condition. (E) The mean PSE in each condition. (F) Mean differences in the PSE between trials preceded by 10% and 90% white-dot stimuli at different lags (1-back, 2-back, and 3-back). * indicates *p* < 0.05; ** indicates *p* < 0.01.

Figure 2B displays the average proportion of “more white dots” responses across the entire experiment. A one-sample t-test conducted to test the null hypothesis that this mean proportion was 0.50 did not yield a significant result (*t*(82) = 0.57, *p* = 0.57), indicating no overall bias in responding toward the white or black category. Figure 2C shows the participants’ accuracy in the perception of proportion across the different actual proportions of white dots. The accuracy increased when the proportion was strongly biased toward either white or black, and decreased as the proportion approached 50%.

Figure 2D shows the accuracy conditioned by the proportion of the preceding trials. Here, the term “condition” refers to the subsets of the dataset divided according to the proportion of white dots in the preceding trial. If the proportion of white dots in the preceding trial was 10%, 40%, 60%, or 90%, the current trial was classified as the Prev10%, Prev40%, Prev60%, or Prev90% condition, respectively. A repeated-measures one-way analysis of variance (ANOVA) comparing accuracy across these four conditions revealed no significant differences (*F*(3, 246) = 1.36, *p* = 0.26). This result suggests that the proportion of the preceding stimulus did not significantly affect the performance or overall perceived difficulty of the current trial.

The mean PSE for each condition are presented in Figure 2E. The plot clearly show that the mean PSE shifted systematically depending on the proportion of white dots in the preceding trial: when the previous proportion was higher, the PSE tended to be higher than when the previous proportion was lower. We first tested whether the mean PSE for each condition matched the mean proportion of “more white dots” responses shown in Figure 2B. One-sample t-tests indicated significant differences for the Prev10% condition (*t*(82) = −3.74, *p* < 0.01) and the Prev40% condition (*t*(82) = −2.64, *p* = 0.01). In contrast, the PSEs in the Prev60% (*t*(82) = −1.00, *p* = 0.32) and Prev90% (*t*(82) = 0.05, *p* = 0.96) conditions were not significantly different from the mean overall response proportion.

Next, a repeated-measures one-way ANOVA was conducted across conditions, which revealed a significant difference in mean PSE between conditions (*F*(3, 246) = 5.77, *p* < 0.01). Post-hoc multiple comparisons using Tukey’s test indicated a significant difference only between the Prev10% and Prev90% conditions (*p* = 0.03), with no other comparisons reaching significance (Prev10% vs Prev40%: *p* = 0.62, Prev10% vs Prev60%: *p* = 0.19, Prev40% vs Prev60%: *p* = 0.86, Prev40% vs Prev90%: *p* = 0.39, Prev60% vs Prev90%: *p* = 0.86). Given that the PSE represents the proportion of white dots perceived as equal, the observed tendency for the PSE to be higher in the Prev90% condition suggests a repulsive effect from the preceding trial, consistent with negative serial dependence. That is, the current perception of proportion was biased in the direction opposite to the proportion of white dots seen in the preceding trial.

To investigate the temporal persistence of serial dependence in the perception of proportion, we conducted an *n*-back analysis. We calculated the difference in the PSE between conditions in which the proportion of white dots *n* trials back was 10% and 90%. The results are shown in Figure 2F. To assess the presence of bias at each *n*-back level (1-, 2-, and 3-back), we conducted one-sample t-tests to determine whether the mean ΔPSE significantly differed from zero. The results revealed that serial dependence was present when the stimulus appeared only from one trial before (1-back), but not for stimuli presented two or more trials earlier (2-back or 3-back) (1-back: *t*(82) = −3.61, *p* < 0.01, 2-back: *t*(82) = 1.02, *p* = 0.31, 3-back: *t*(82) = −1.06, *p* = 0.29). Furthermore, the sign of ΔPSE reflects the direction of the serial dependence effect. A positive value indicates an attractive bias, meaning that perceptual choices are drawn toward the previous stimulus. In contrast, a negative value indicates a repulsive bias, in which choices are shifted away from the previous stimulus. The serial dependence observed in the 1-back condition as found to be repulsive, as indicated by the negative value of the ΔPSE. Therefore, in the 1-back condition, the serial dependence between the Prev10% and Prev90% conditions showed the same pattern in the analysis presented in Figure 2F as in Figure 2E.

### 3.2 Experiment 2 (Dot vs Diamond)

In Experiment 2, as in Experiment 1, we first assessed whether the observed effects could be explained by non-perceptual response patterns such. The proportion of trials in which a participant’s current response repeated their previous response was analyzed. The results, shown in Figure 3A, indicated no systematic tendency toward either repetition or alternation for either the right key (*t*(77) = 0.57, *p* = 0.57) or the left key (*t*(77) = 1.73, *p* = 0.09). This suggests that the observed serial dependence effects are perceptual and not artifacts of unconscious response strategies.

**Figure 3:**
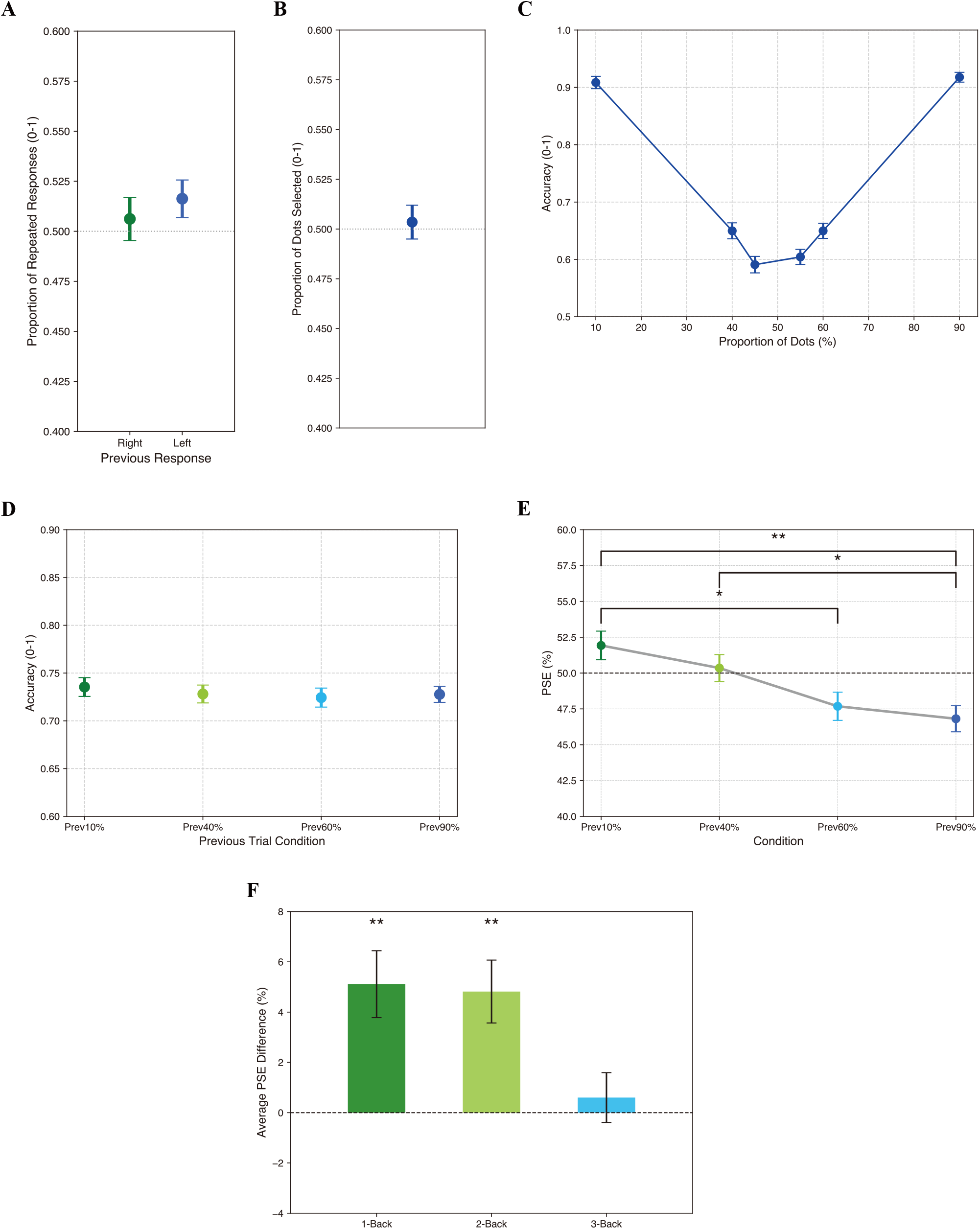
Results from Experiment 2. Error bars in (A)–(D), and (F) represent the standard error of the mean across participants. (A) Probability of repeating the same response as the previous trial. (B) The mean proportion of “more dots” responses across all participants. (C) Overall accuracy across participants for each proportion of dots. (D) Overall accuracy across participants for each condition. (E) The mean PSE in each condition. (F) Mean differences in the PSE between trials preceded by 10% and 90% dot stimuli at different lags (1-back, 2-back, and 3-back). * indicates *p* < 0.05; ** indicates *p* < 0.01.

The overall mean proportion of “more dots” responses across the entire experiment is displayed in Figure 3B. A one-sample t-test conducted against the null hypothesis of a 0.50 mean proportion was not rejected (*t*(77) = 0.41, *p* = 0.69), confirming no overall response bias toward the “dots” or “diamonds” category. Figure 3C illustrates the relationship between actual dot proportion and accuracy. Consistent with Experiment 1, accuracy followed the expected pattern: it decreased as the proportion of dots approached 50% and increased for more extreme (skewed) proportions.

To evaluate whether the preceding trial’s proportion affected the difficulty of the current task, a repeated-measures one-way ANOVA on accuracy across the four preceding proportion conditions was performed (Figure 3D). The analysis revealed no significant differences in accuracy across the Prev10%, Prev40%, Prev60%, and Prev90% conditions (*F*(3, 231) = 0.45, *p* = 0.72). This indicates that the proportion of the previous stimulus did not influence the overall task performance or perceived difficulty of the current trial in Experiment 2.

Figure 3E presents the mean PSE across participants for each preceding proportion condition. The plots show that, contrary to Experiment 1, the PSE generally shifted toward the preceding proportion. Specifically, when the previous proportion was lower (e.g., Prev10%), the current PSE tended to be higher, and when the previous proportion was higher (e.g., Prev90%), the current PSE tended to be lower. We tested the null hypothesis that the mean PSE for each condition was equal to the mean proportion of “more dots” responses. Separate one-sample t-tests revealed significant differences for the Prev60% (*t*(77) = −2.69, *p* = 0.01)) and Prev90% (*t*(77) = −3.85, *p* < 0.01) conditions, but no significant differences for the Prev10% (*t*(77) = 1.57, *p* = 0.12) and Prev40% (*t*(77) < 0.01, *p* = 1.00) conditions.

A subsequent repeated-measures one-way ANOVA across the four conditions confirmed a significant effect of the preceding proportion on the PSE (*F*(3, 231) = 9.26, *p* < 0.01). Posthoc Tukey’s tests indicated significant differences between the Prev10% and Prev60% conditions, the Prev10% and Prev90% conditions, and the Prev40% and Prev90% conditions (Prev10% vs Prev40%: *p* = 0.66, Prev10% vs Prev60%: *p* = 0.01, Prev10% vs Prev90%: *p* < 0.01, Prev40% vs Prev60%: *p* = 0.21, Prev40% vs Prev90%: *p* = 0.05, Prev60% vs Prev90%: *p* = 0.92). This systematic shift demonstrates positive serial dependence (attractive bias). This means that a low proportion of dots in the preceding stimulus biased the current perception toward a lower proportion than its actual value, and conversely, a high preceding proportion biased the current perception toward a higher proportion.

Finally, the *n*-back analysis (Figure 3F) was performed to examine the temporal persistence of serial dependence in perception of proportion. One-sample t-tests on the PSE difference (ΔPSE) between the 10% and 90% proportion conditions revealed that serial dependence was significantly present for stimuli presented 1-back (*t*(77) = 3.84, *p* < 0.01) and 2-back (*t*(77) = 3.85, *p* < 0.01), but not for stimuli presented 3 trials earlier (3-back: *t*(77) = 0.61, *p* = 0.55). Furthermore, the direction of the ΔPSE in the 1-back condition was consistent with the attractive bias (positive serial dependence) found in the main analysis (Figure 3E).

## 4 Discussion

Serial dependence, in which previous visual input systematically biases current perception, has been established across a range of features, including low-level attributes like orientation [6–10] and color [14–16]. However, its generalization is not yet well understood. The present study was designed to investigate whether serial dependence also emerges in the perception of proportion, a previously unexplored visual feature. The results demonstrated that serial dependence emerges in proportion judgments, but critically, the direction of the bias was not consistent and varied depending on the features defining the proportion. Furthermore, these findings extend previous research by investigating the influence of response confidence and stimulus similarity on this novel form of serial dependence.

We observed a repulsive serial dependence effect in Experiment 1 and an attractive serial dependence effect in Experiment 2. In Experiment 1, where proportion was defined by brightness (white vs. black dots), the finding of a repulsive effect demonstrates that serial dependence arises in brightness-based proportion judgments. Nevertheless, this design left open the possibility that the observed serial dependence was specific to the brightness attribute of the stimuli. To address this generalization, Experiment 2 employed two types of symbols, dots and diamonds, which differed only in shape but were equated for brightness. The results showed a significant attractive effect of the preceding stimulus, indicating that serial dependence is a general phenomenon that also emerges in shape-based proportion judgments. This highlights that serial dependence in proportion is robust across different feature domains.

The influence of serial dependence was found to diminish over time in the present study. The present study demonstrated that serial dependence extends up to two trials back. This finding is consistent with previous research. Fischer and Whitney [5], for example, observed that while the strength of serial dependence in orientation judgments decreased over time, the effect could persist up to three trials back. The longer inter-trial intervals likely inherent in their adjustment task, where participants manually rotated a response bar, may have contributed to this extended temporal persistence. However, studies employing two-alternative forced choice (2AFC) tasks―like the present study―have reported a similar temporal decay in serial dependence [35]. Consistent with these findings, both Experiment 1 (Brightness Proportion) and Experiment 2 (Shape Proportion) in the present study revealed this temporal decay. Notably, in Experiment 2, where the overall magnitude of serial dependence was greater, the effect persisted up to two trials back.

Confidence in response in the preceding trial may influence subsequent serial dependence. Samaha et al. [36] investigated how participants’ confidence influenced serial dependence by having them report both their responses to the task and their confidence in those responses. The results showed that serial dependence became stronger when participants had higher confidence in the preceding trial, and weaker when their confidence was lower. In the present experiment, when the test stimulus consisted of white and black symbols in Experiment 1, serial dependence was not observed for the pair of trials in which the proportion in the immediately preceding trial was 40% and 60%. These findings extend the work of Samaha et al. Although participants in the current study did not explicitly report their confidence, an inference can be drawn from the accuracy rates across different proportions of white dots (Figure 2C and Figure 3C). Specifically, it is likely that participants experienced greater judgmental confidence following trials that contained a highly imbalanced ratio of the two shape types. Conversely, confidence was likely attenuated following trials where the preceding stimulus was more balanced. This lack of confidence might have diminished the influence of the previous stimulus. Such a reduction could explain why serial dependence was not observed for the pairs where the proportion on the immediately preceding trial was 40% or 60%.

Serial dependence may be influenced by participants’ attention. In Experiment 1, the absence of serial dependence between the Prev40% and Prev60% conditions was attributed to participants’ confidence in their responses. If confidence were the only factor influencing serial dependence, then the gap of biases between the Prev40% and Prev60% conditions should also have been absent in Experiment 2, the accuracy in which was lower than in Experiment 1. However, Experiment 2 revealed a significant attractive serial dependence overall. This suggests that the influence of attention may have played a significant role. Fischer and Whitney [5] demonstrated that attention to stimulus features can modulate the magnitude of serial dependence. In the present study, the stimuli in Experiment 2 were composed exclusively of white symbols. As indicated by the accuracy results shown in Figure 2C and Figure 3C, this made it more difficult for participants to discriminate proportions compared to Experiment 1. This increased difficulty likely required participants to allocate greater attention to the stimuli in Experiment 2. This enhanced attentional engagement could have amplified the serial dependence effect, leading to its emergence in Experiment 2 but not in Experiment 1.

The opposing serial dependence effects―repulsive (Experiment 1, Brightness) and attractive (Experiment 2, Shape)―are likely attributable to stimulus similarity within the feature space. Prior work by Fritsche and de Lange [37] demonstrated that how similarity between successive stimuli affects serial dependence. When successive stimuli were similar, responses were biased toward the previous stimulus (attractive bias), whereas larger differences led to repulsive biases. This suggests that the proximity between stimuli in the underlying feature space is a critical determinant of the bias direction. In the present study, comparing the results of Experiments 1 and 2 revealed opposite patterns of serial dependence. This difference may reflect the role of feature space proximity, as indicated in prior studies. Experiment 1 used white and black symbols, introducing a significant brightness contrast across the display. In contrast, Experiment 2 used only white symbols (dots and diamonds). Consequently, even for an equivalent magnitude of proportional change between trials, the presence of varied brightness in Experiment 1 likely resulted in a greater perceived difference between successive stimuli in the relevant feature space (brightness/contrast), pushing the bias toward repulsion (negative serial dependence). Conversely, the high similarity and low contrast in Experiment 2 likely kept the stimuli closer in feature space, leading to attraction (positive serial dependence).

Serial dependence may originate from higher-level cognitive processes. Previous studies have typically used stimulus features such as direction [6–10], color [14–16], or spatial position [11–13] presented in the immediately preceding trial. In such experiments, participants were only required to reproduce what they had just seen, without the need for detailed processing of the stimulus content. As a result, serial dependence has often been regarded as a phenomenon that arises primarily at the low-level sensory stage [5, 6, 12]. However, several studies have shown that post-perceptual processes such as working memory and decision-making also contribute to serial dependence [8, 37–39]. These discrepancies have led to ongoing debate over whether serial dependence arises from low-level sensory processing or higher-level cognitive processing [8, 10, 11]. In the present study, serial dependence was examined in the context of “proportion”, where participants were required to judge which of two presented symbols contains a greater proportion. This task involved not merely reproducing perceived information but a series of processes, including integrating and comparing information before making a decision and responding. Thus, compared to previous studies that focused on low-level perceptual features, this task demands a greater degree of higher-order cognitive processing. These findings suggest that serial dependence may arise from high-level processing. This interpretation aligns with existing research that has investigated the involvement of higher-level mechanisms in serial dependence [40].

This study successfully demonstrated the presence of serial dependence in the perception of proportion judgments; however, it has several limitations. One major limitation is that it remains challenging to fully distinguish proportion-based serial dependence from numerosity-based serial dependence. Fornaciai and Park [20] examined serial dependence in numerical judgments using dot arrays with equal proportions of white and black dots. In each trial, participants viewed a sequence of stimuli and judged whether the final array contained more or fewer dots than a preceding one. In this case, the dots were not distinguished by brightness and were treated as a single stimulus composed of two brightness levels. The study found attractive serial dependence in numerical judgments. In contrast to these previous studies, our experiment manipulated the relative proportion of white and black dots (Experiment 1), or dots and diamonds (Experiment 2), thereby treating the two symbols as two kinds of stimulus. Furthermore, by shifting from brightness-based to shape-based proportion judgments, we increased the difficulty of individuation and reduced the influence of numerical judgments. However, in both experiments, participants could still potentially count the number of symbols. Therefore, we cannot completely rule out the possibility that numerosity-based serial dependence influenced their responses. Future studies should consider developing methods that manipulate stimulus proportion while keeping the total number of stimuli constant. Such approaches would help disentangle proportion-based and number-based serial dependence.

Moreover, a limitation of the present study is the inability to quantitatively assess stimulus similarity across trials. We interpreted the shift from repulsive bias (Experiment 1, Brightness) to attractive bias (Experiment 2, Shape) as reflecting the degree of similarity between consecutive stimuli, with higher similarity leading to attraction. While the brightness contrast in Experiment 1 intuitively suggests a greater feature distance compared to the shape difference in Experiment 2, this similarity is difficult to quantify precisely. In studies using orientation, similarity is easily captured by the angular difference between stimuli [37]. However, in our study, the dot arrays were randomly distributed spatially on each trial. Although the proportional difference is quantifiable, the unquantified differences in the spatial arrangement and perceived feature distance contribute to the characteristics of serial dependence. This difficulty in assigning a simple numerical scale to the perceptual similarity of complex, spatially variable dot arrays highlights the need for future methodological work to systematically quantify perceptual similarity in such stimuli.

Additionally, the online experimental environment presents inherent limitations. As the study was conducted via an online platform rather than a controlled laboratory setting, we could not employ direct monitoring methods like eye-tracking to ensure gaze fixation or sustained attention. Although we implemented screening based on no-response rates and accuracy to exclude inattentive participants, and imposed a response time limit to promote quick engagement, we cannot assess participants’ attentional state on a trial-by-trial basis. Consequently, the influence of unobserved attentional lapses on trial-level responses cannot be entirely ruled out.

Finally, the ecological validity of the experimental setup remains limited. Our study presented only two types of symbols on a uniform gray background. In contrast, real-world visual environments are saturated with numerous stimuli possessing a variety of features (e.g., physical distance, weight) beyond the target of attention. The presence and functional significance of serial dependence under such more complex and naturalistic conditions are still largely unknown. While this study provides fundamental insights, future research should aim to develop experimental settings that more closely approximate the richness of real-world environments to fully understand the ubiquity of serial dependence.

To conclude, the present study investigated the behavioral characteristics of serial dependence in the perception of proportion. These findings contribute to a more comprehensive understanding of the effect and offer new evidence relevant to the ongoing debate concerning the nature of serial dependence. Our results demonstrate that past perception of proportion biases the perception of current proportions. This bias, consistent with previous research, is modulated by factors such as participant confidence and the similarity between successive stimuli in feature space.

## Data and code availability

The experimental data and analysis code used in this study are publicly available on GitHub: https://github.com/HKaede/SD_proportion_perception.

## Acknowledgments

This work is supported in part by JSPS KAKENHI with grant number 23K26110, 23K17461, 22H05163, and 24K15047, and JST AdCORP under Grant JPMJKB2307.

The Gemini 2.5 Flash generative AI tool (Source: Google) was utilized for proofreading of the English manuscript on October 20, 2025.

